# Protein structure prediction using AI and quantum computers

**DOI:** 10.1101/2021.05.22.445242

**Authors:** Ben Geoffrey A S

## Abstract

This work seeks to combine the combined advantage of leveraging these emerging areas of Artificial Intelligence and quantum computing in applying it to solve the specific biological problem of protein structure prediction using Quantum Machine Learning algorithms. The CASP dataset from ProteinNet was downloaded which is a standardized data set for machine learning of protein structure. Its large and standardized dataset of PDB entries contains the coordinates of the backbone atoms, corresponding to the sequential chain of N, C_alpha, and C’ atoms. This dataset was used to train a quantum-classical hybrid Keras deep neural network model to predict the structure of the proteins. To visually qualify the quality of the predicted versus the actual protein structure, protein contact maps were generated with the experimental and predicted protein structure data and qualified. Therefore this model is recommended for the use of protein structure prediction using AI leveraging the power of quantum computers.

## Introduction

The past few years have seen a surge in the application of data-driven AI methods to solve complex problems in biology [1-3]. The areas of application include genomics, proteomics, and drug discovery[4-10]. These have led to tremendous advances in research in biological science. Another emerging area from which life science research can leverage advances is through quantum computing[11-15]. Therefore this work seeks to combine the combined advantage of leveraging these emerging areas of Artificial Intelligence and quantum computing in applying it to solve the specific biological problem of protein structure prediction using Quantum Machine Learning algorithms. Quantum machine learning algorithms are machine/deep learning algorithms that leverage the power of quantum computers. Fault-tolerant quantum computers may be far off, however solving real-world quantum-chemistry problems on near-term quantum devices is possible through the Pennylane [16] which provides an interface to use any of the quantum hardware provided by any of the quantum hardware providers such as IBM, Google, or Microsoft. Classical data is encoded into quantum states through amplitude encoding techniques to reflect the quantum state/wave-function of the quantum circuit. The circuit parameters are then optimized to produce the solution. These quantum layers are introduced between classical Keras layers to turn classical Keras-based deep neural network into a quantum-classical hybrid deep neural network to solve data-driven AI problems through leveraging the power of quantum computers.

## Methods

The CASP dataset from ProteinNet was downloaded. ProteinNet is a standardized data set for machine learning of protein structure [17]. Its large and standardized dataset of PDB entries contains the coordinates of the backbone atoms, corresponding to the sequential chain of N, C_alpha, and C’ atoms. This dataset was used to train a quantum-classical hybrid Keras deep neural network model whose model configuration is shown below in Fig. 1. A classical convolutional deep neural network inspired in part by Wang et. al. [18] along with introducing quantum Keras layers to make a hybrid model was built and the summary of input dimension, hidden layers, and output dimension of the network is shown in Fig.1.

**Fig. 1.**
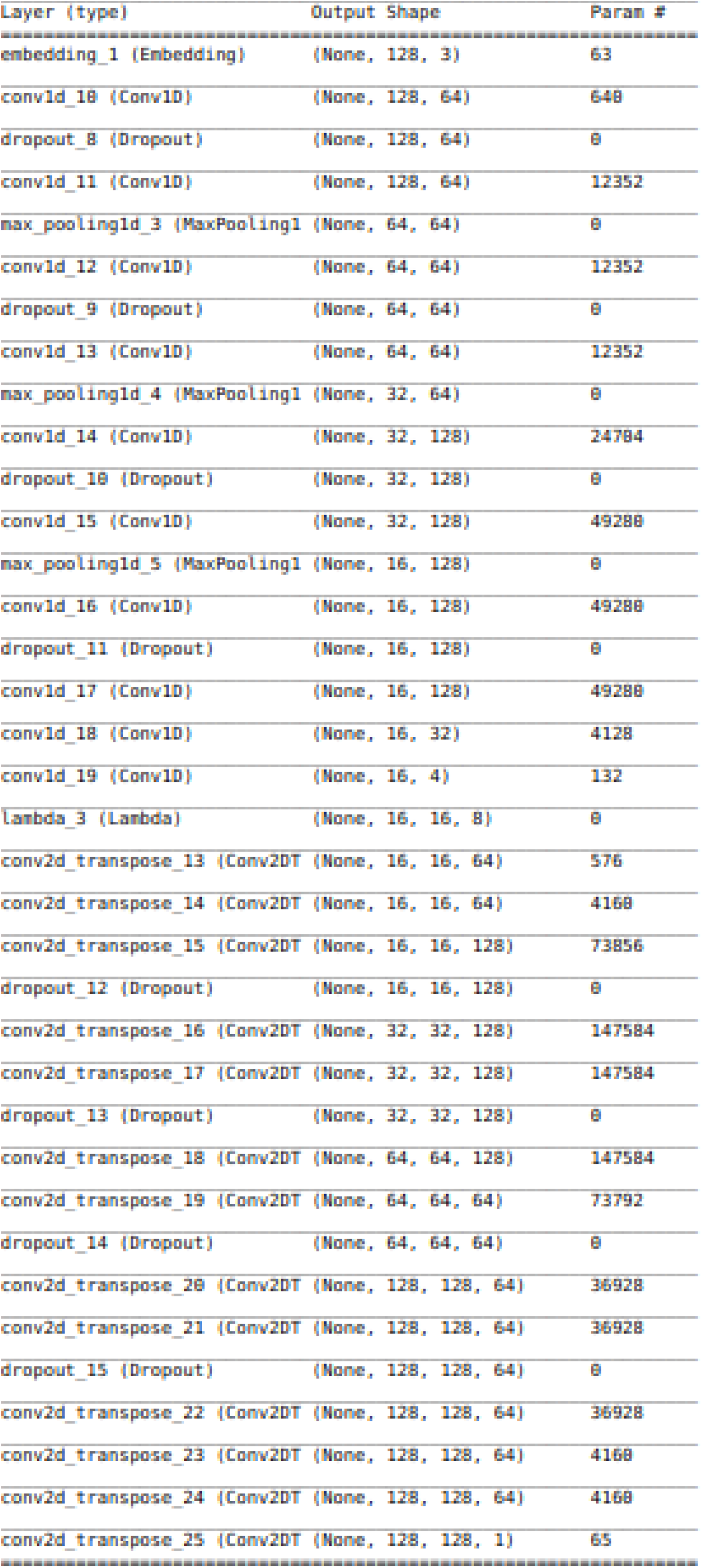
Model Configuration

## Results and Discussion

With the trained model, predictions were carried out. To visually qualify the quality of the predicted versus the actual protein structure, protein contact maps were generated with the actual and predicted protein structure data. The predicted and the actual contact map for two instances of prediction for a random seed is shown below in Fig.2

**Fig. 2.**
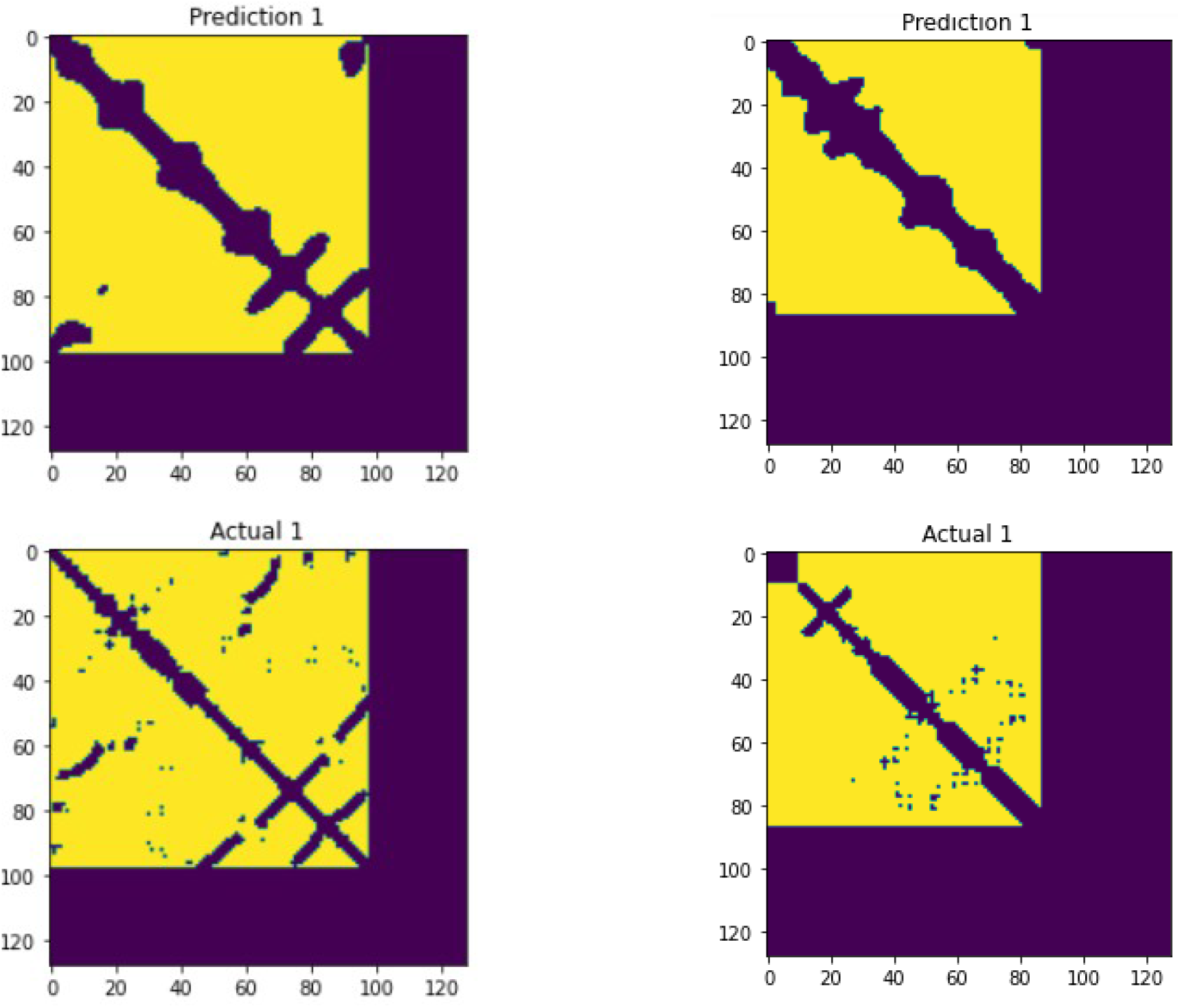
Protein contact maps

The converegnce of the model is depicted in the figure below

**Fig. 3.**
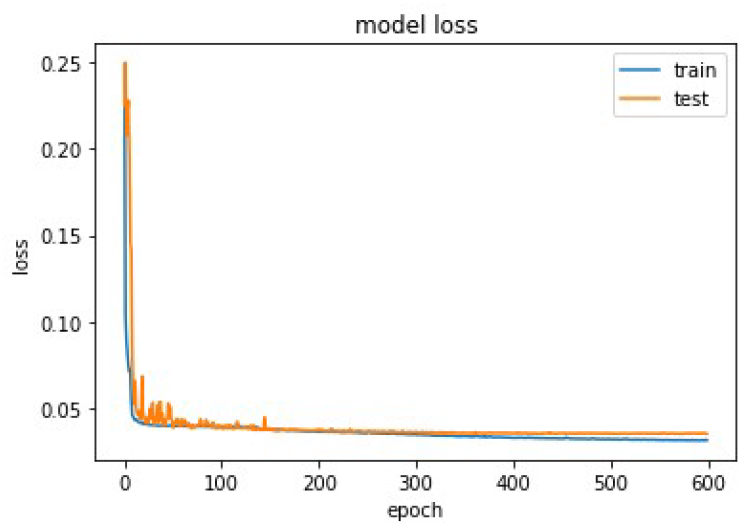
Loss function convergence

